# Modulation of Genome Editing Outcomes by Cell Cycle Control of Cas9 Expression

**DOI:** 10.1101/127068

**Authors:** Yuping Huang, Caitlin McCann, Andrey Samsonov, Dmitry Malkov, Gregory D Davis, Qingzhou Ji

## Abstract

Targeting specific chromosomal sequences for genome modification or regulation during particular phases of the cell cycle may prove useful in creating more precise, predictable genetic changes. Here, we present a system using a fusion protein comprised of a programmable DNA modification protein, Cas9, linked to a cell cycle regulated protein, geminin, as well as green fluorescent protein (GFP) for visualization. Despite the large size of Cas9 relative to geminin, cells were observed to express Cas9-GFP-geminin at levels which oscillate with the cell cycle. These fusion proteins are also shown to retain double-strand break (DSB) activity at specific chromosomal sequences to produce both indels and targeted integration of donor ssDNA. Most importantly, the ratio of ssDNA donor integration to non-homologous end joining (NHEJ) was observed to increase, suggesting that cell cycle control Cas9 expression may be an effective strategy to bias DNA repair outcomes.

## 1. Introduction

Genome editing with targeted nucleases has gained wide adoption for gene knockout, disease single nucleotide polymorphism (SNP) modeling, and tagging genes within their endogenous chromosomal contexts with reporter proteins. The key molecular technology enabling this revolution in genome editing arose from the discovery that targeted double strand breaks (DSBs) caused thousand-fold increases in DNA repair rates in mammalian cells[1]. One revolutionary, increasingly popular method to create such DSBs is the CRISPR system. This system is naturally present in various species of bacteria and archaea, where it functions as an adaptable immune system. Recently, it has been exploited as a system for genome editing[2,3]. In this system, CRISPR-associated (Cas) nucleases are guided to a target sequence by a complementary guide RNA, where they introduce a DSB with high specificity. CRISPR offers advantages over other genome editing systems such as zinc-finger nucleases (ZFNs) and transcription activator-like effector nucleases (TALENs), primarily because of its RNA-guided nature enabling facile reprogramming to nearly any target sequence of interest. Designing a CRISPR guide RNA is much easier and more affordable than engineering a new ZFN or TALEN protein to target a new sequence, and the target site has no sequence limitation except that the sequence is bordered by a short protospacer adjacent motif (PAM)[2,3].

DSBs, such as those introduced by Cas9, can be repaired via two primary pathways in mammalian cells: non-homologous end-joining (NHEJ) or homology-directed repair (HDR). The NHEJ pathway is error-prone, which leads to mutations in genes at the target sites through the creation of insertions and deletions (indels); therefore, this repair pathway is effective in creating gene knockouts and loss-of-function alleles, but the HDR pathway is needed for applications requiring more precise editing or the insertion of exogenous synthetic sequences[4]. While NHEJ is highly efficient in many cell types and relatively easy to attain, achieving high rates of HDR has proven more challenging[5–7].

The HDR pathway precisely repairs DNA damage through copying a second, homologous ‘donor’ dsDNA into the damaged site. If the dsDNA donor sequence exactly matches the sequence of the damaged site before the breakage or insult occurred, then the repair precisely restores the DNA to its undamaged state[8]. Alternatively, a single-stranded oligonucleotide donor (ssODN) can effectively integrate a new sequence at the damaged site by processes which are suspected to resemble classical HDR for dsDNA donors[7]. This can be exploited to create gene knock-ins, integrate transcriptional control sequences or RNA coding sequences, and more[5–7].

It is hypothesized that HDR may have evolved in mammalian cells to more accurately repair damage that occurs specifically during DNA replication, as the HDR pathway is cell cycle dependent. The cell cycle-dependent nature of HDR is governed by many factors. For example, transcription of HDR genes varies throughout the cycle and the mechanism itself requires sister chromatids. In addition, many HDR proteins must be activated through phosphorylation by cyclin-dependent kinases (CDKs), which regulate the cell cycle and are themselves regulated by the cycle[8].

The mechanism of the cell cycle is intricate and versatile. Progression through the cell cycle is controlled by both intracellular and extracellular signals in a series of complex, intertwined pathways with feedback and feed-forward mechanisms. This complexity allows the cycle to be coordinated with differentiation, morphogenesis, apoptosis, and more[9]. Cell cycle phase completion times are in many ways phenotypes of a given cell, varying by cell type, cellular environment, and metabolic stage[10]. For example, human embryonic stem cells are observed to have a dramatically shortened G1 phase[11], which lengthens as the cells differentiate into their final cell types[12]. Even within a cell type, individual cells can have variability in cell cycle phases[13]. Because HDR is found only in the S and G2 phases of the cell cycle, differences in cell cycle phases can have a significant potential impact on the outcomes of genome editing experiments. Thus, for certain genome editing scenarios, it is advantageous to limit DNA damage to only these phases.

The cell cycle has many points of regulation that can be exploited for the purpose of limiting genome editing to particular phases of the cell cycle. Chemical or genetic disturbances to the NHEJ pathway can promote the HDR pathway, but these are often difficult to implement and may prove too harmful to cells to create a working model[14]. Transcription can be up-or down-regulated through the manipulation of promoters, inhibitors, transcription factors, and more. MicroRNAs can cause degradation of mRNAs that encode regulatory proteins before they are translated into functional proteins[15]. Unfortunately, control at either the transcriptional or post-transcriptional levels regulates when proteins are expressed, but fails to govern when proteins are degraded; in other words, proteins expressed in one phase may carry into the next. For this reason, we favored a post-translational method of cell cycle regulation. Cell cycle regulation can occur at the post-translational level through a process called ubiquitination, in which the addition of one or more ubiquitin (Ub) molecules to a protein regulates the protein’s activity or leads to its degradation by the proteasome[9].

To encourage Cas9 activity during the S and G2 phases, we searched for fusion protein candidates that are targeted for degradation during the M phase and/or early G1 phase of the cell cycle. One such protein is geminin, which is a substrate of the APC/Chd1 complex, the primary cell cycle-controlling E3 ubiquitin ligase. Human geminin is a DNA replication inhibitor approximately 25 kDa in size. It is expressed during the S and G2 phases of the cell cycle and is inactivated through Ub-mediated proteolysis during the metaphase-anaphase transition[9]. Therefore, geminin is only active in the S, G2, and M phases.

Here, we present a novel approach to increase HDR via the fusion of Cas9 to GFP and the first 110 amino acids of geminin, creating a programmable, cell cycle-regulated DNA modification protein. This system allows expression of Cas9 in a cell cycle dependent manner, was observed to increases ratios of ssDNA oligo integration rates relative to indels.

## 2. Materials and Methods

### 2.1 CRISPR Design and Construction

The Cas9-GFP-geminin plasmid was generated by subcloning to produce a recombinant Cas9-GFP-geminin fusion protein. The backbone blast is Lv5-EF1α-Cas9-Blast (MilliporeSigma, Lot# LVCAS9BST), which contains the unique XbaI (NEB, R0145T) and HpaI (NEB, R0105S) restriction enzyme sites used in cloning. The geminin gene inserted was carried by a pUC57 plasmid vector from GenScript (product# 648334-1). This gene was cut from the parent plasmid using XbaI (NEB, R0145T) and HpaI (NEB, R0105S) restriction enzymes. The backbone was cut using the same enzymes, creating compatible ends. The geminin insert was then ligated into the backbone to create a Cas9-GFP-geminin fusion protein expression vector. The Cas9 only plasmid used was a product of Cas9GFP (MilliporeSigma, Lot# 05271523MN), which uses a CMV promoter for strong, transient expression of Cas9.

The *AAVS1*-sgRNA used was from pU6-gRNA (MilliporeSigma, Lot#05271523MN). The single-stranded oligonucleotide donor (*AAVS1*-ssODN) was designed such that successful integration would generate a unique SpeI (NEB, R0133S) restriction enzyme site. In the sequences below, this guanine to thymine mutation is seen in red, and the SpeI site is underlined. This oligonucleotide donor was custom synthesized through MilliporeSigma.

AAVSI Target Sequence

5’- GGGCC*<u>ACTAG**G**</u>*GACAGGATTGG -3’

Integrated *AAVS1* Oligonucleotide Donor with Spel Mutation

5’-GTTCTGGGTACTTTTATCTGTCCCCTCCACCCCACAGTGGGGCC<u>ACTAG**T**</u>GACA GGATTGGTGACAGAAAAGCCCCATCCTTAGGCCTCCTCCTTCCTAG -3’

### 2.2 Cell Lines and Transfection

The human cell lines K562 and U2-OS were acquired from ATCC. K562 cells were maintained in DEME media (cat# I3390, MilliporeSigma) supplemented with 10% fetal bovine serum and 2 mM L-Glutamine. U2-OS cells were grown in McCoy’s 5A media (cat# M8403, MilliporeSigma) supplemented with 10% fetal bovine serum and 1.5 mM L-Glutamine; all supplements were obtained from MilliporeSigma. Cells were kept in a 37°C, 5% CO_2_ incubator.

The cells were consistently split 1 day prior to transfecting and transfections were performed at ∼80% confluency for U2-OS cells, or ∼1 million cells per ml for K562 cells. The cells were nucleofected using an Amaxa^®^ Nucleofector® with Amaxa® Cell Line Nucleofector® Kit V (Lonza) using the manufacturer’s protocol. Both the U2-OS cells and K562 cells were suspended in Nucleofector™ Solution V at a concentration of 1x10^6^ cells/100 μl with 4 μg of Cas9 and each gRNA construct and 3 μl of 100 μΜ ssODNs in a 4mm gap cuvette (BioExpress, Kaysville, UT). The K562 cells were nucleofected using program T-016, while the U2-OS cells were nucleofected using program X-001. Cells were then plated on 6-well plates with 2 ml complete growth media per well at 37°C for 48 hours prior to analysis.

### 2.3 CEL-1 Analysis of CRISPR/Cas9 Cleavage Activity

SURVEYOR, a type of CEL enzyme, is a mismatch-specific nuclease used to detect single-base mismatches or small insertions or deletions. This enzyme is capable of cleaving at multiple mutations in large DNA fragments, resulting in cleavage products that can then be detected via gel electrophoresis. Genomic DNA was isolated from the transfected cells using a GenElute™ Mammalian Genomic DNA Miniprep kit (MilliporeSigma, G1N70). This was done following the manufacturer’s protocol, but the DNA was eluted with 200 μl of sterile water (W4502), rather than the provided elution buffer.

The genomic region surrounding each mutant target gene locus was amplified via PCR using JumpStart™ Taq ReadyMix (MilliporeSigma, P2893) under the following conditions: 98°C for 2 min; 35x (98°C for 15 s, 62°C for 30 s, 72°C for 45 s); 72°C for 5 min; hold at 4°C. The *AAVS1* primers used for PCR are supplied in Table 1. Then, 200 ng of each PCR product was subjected to heteroduplex formation via the following annealing program: 95°C for 10 minutes, 95°C to 85°C with a ramp rate of −2°C/s, 85°C to 25°C with a ramp rate of −0.1 °C/s, hold at 4°C. The heteroduplexed products were treated with SURVEYOR Nuclease S and SURVEYOR Enhancer S (IDT Technologies) for 30 minutes at 42°C, then samples were electrophoresed by running them on a nondenaturing 10% TBE PAGE gel (BioRad). The gel was stained in 100 ml 1xTBE buffer with 3 μl of 10 mg/ml ethidium bromide for 5 min, then destained with 1 xTBE buffer and visualized with a UV illuminator. The resulting bands were quantified using Image J software, and percent cutting was calculated using the following formula:

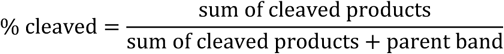

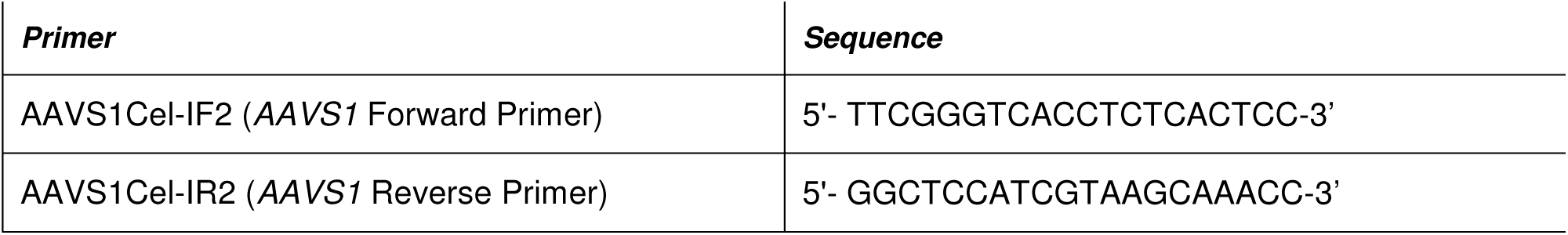
Table 1:

### 2.4 RFLP Analysis of CRISPR/Cas9 Cleavage Activity

Restriction Fragment Length Polymorphism (RFLP) analysis is a technique used to detect variations in homologous DNA sequences. In RFLP analysis, the DNA sample is broken into pieces and digested by restriction enzymes, then the resulting restriction fragments are separated according to their lengths via gel electrophoresis. The same PCR products subjected to SURVEYOR assay, above, were also used in this RFLP analysis. For this assay, the PCR products were purified using a GenElute™ PCR Clean-Up Kit (MilliporeSigma, NA1020). Then, 200 ng of each PCR product was digested with SpeI restriction enzyme (NEB, R0133S) following the manufacturer’s protocol. The digested DNA samples were then separated on an ethidium bromide-stained 10% acrylamide TBE gel (BioRad), visualized with a UV illuminator, and quantified using Image J software.

### 2.5 Imaging and Image Analysis

Fluorescent imaging of cells was performed using a Nikon Eclipse TE2000-E inverted research microscope, a Photometrics CoolSNAP ES^2^ cooled CCD camera, and MetaMorph® software using a 20x/0.75 air objective. Cells were imaged live in Hanks Balanced Salt Solution (H8264) supplemented with 2% fetal bovine serum (F2442). The filterset was GFP (ex 450–490/em 500–550). The localization and redistribution of Cas9-geminin-GFP fusion protein during 24 hours was done in a full enclosure stage top incubator for live microscopy (In Vivo Scientific, USA).

## 3. Results and Discussion

HDR is restricted to the S and G2 phases of the cell cycle and coordinating the cell cycle and the delivery of Cas9 ribonucleoproteins has been shown to increase HDR-facilitated genome editing[16]. Previous experiments to synchronize cells or enhance HDR for genome editing by small molecule cell cycle regulators, singlestranded DNA oligonucleotides, and chemical NHEJ inhibitors have shown limitations such as cytotoxicity, cell-type dependence, and a limited increase in HDR[7,16,17]. Inspired by the visualization of a cell cycle dependent geminin-GFP fusion created to study cell cycle dynamics[9], we created a new protein by fusing Cas9 to GFP and the N-terminal of the first 110 amino acids of geminin. In doing so, we hope to create a GFP-visualized Cas9 protein that is targeted for proteasomal degradation in late mitosis or early G1, synchronizing its activity to the phases of the cycle in which HDR is present.

### 3.1 Cas9-GFP-Gemimin fusion protein expression is cell cycle dependent

To construct the Cas9-GFP-geminin fusion protein, we fused a fragment encoding the first 110 amino acid residues of geminin into an Lv5-EF1α-Cas9-Blast backbone. This produced a Cas9-GFP-geminin fusion protein expression vector (Figure 1) with Cas9 on the N-terminus of a GFP-geminin fusion. The DNA and amino acid sequences for this fusion protein can be found in supplementary Tables S1 and S2, respectively. Because the molecular weight of SpCas9 is approximately 10-fold larger than that of geminin, we needed to ensure that ubiquitin-mediated functions were maintained sufficiently to degrade our fusion protein in an S/G2-specific manner. Upon transfecting the Cas9-GFP-geminin plasmid into U2-OS cells, visualization of the cells via fluorescence microscopy confirmed the expression of Cas9-GFP-geminin. The GFP signal was observed to slowly increase throughout the S and G2 phases, and then rapidly decrease below detection just before cell division in the G2 or early M phases, reappearing in the two daughter cells in the S phase, suggesting that the cycling functions of geminin remained intact when fused to Cas9. The cyclic dynamics of the fusion protein can be seen in Figure 2A, which includes snapshots taken from a video (Supplementary Video 2). The cell cycle dependent expression of the fusion protein is graphed in Figure 2B.

**Figure 1.**
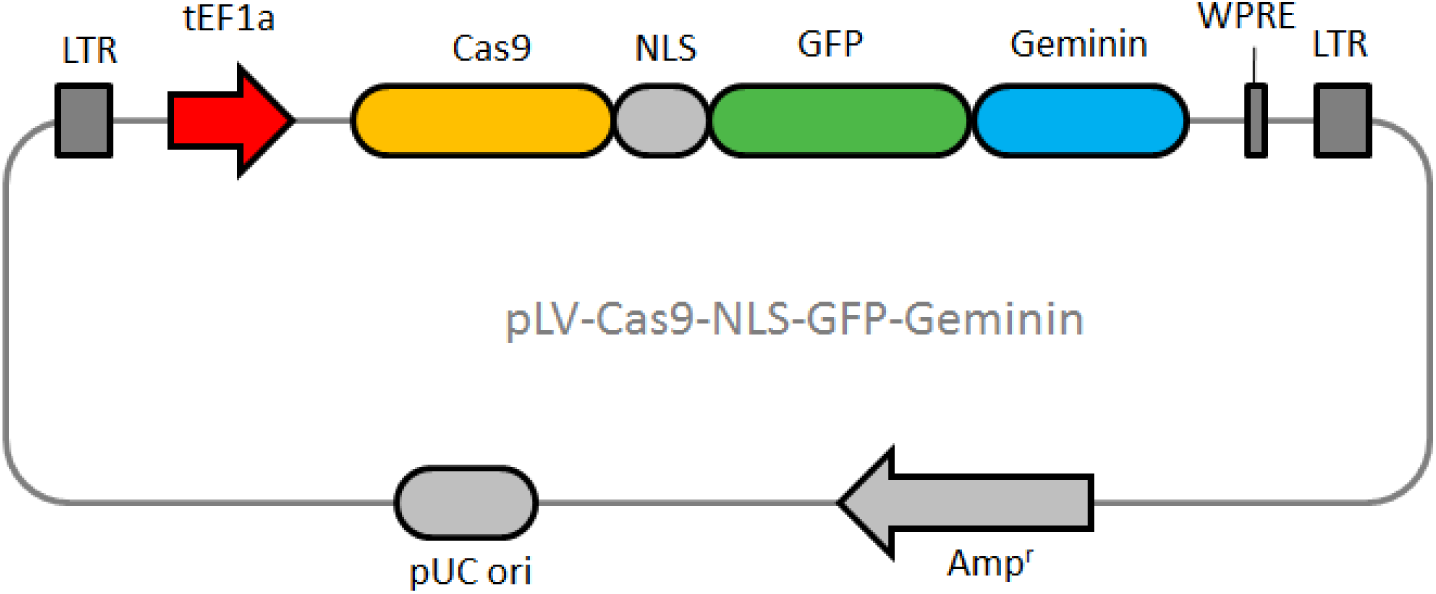
pLV-Cas9-GFP-geminin Plasmid Map. The Cas9-GFP-geminin plasmid was generated by subcloning genetic sequence for the first 110 amino acid residues of geminin into in the Lv5-EF1α-Cas9-Blast backbone. LTR, long terminal repeats; tEF1α, elongation factor 1-alpha; NLS, nuclear localization signal; GFP, green fluorescent protein; WPRE, Woodchuck Hepatitis Virus (WHP) Posttranscriptional Regulatory Element; pUC ori, origin of replication from pUC; Amp^r^, ampicillin resistance.

**Figure 2.**
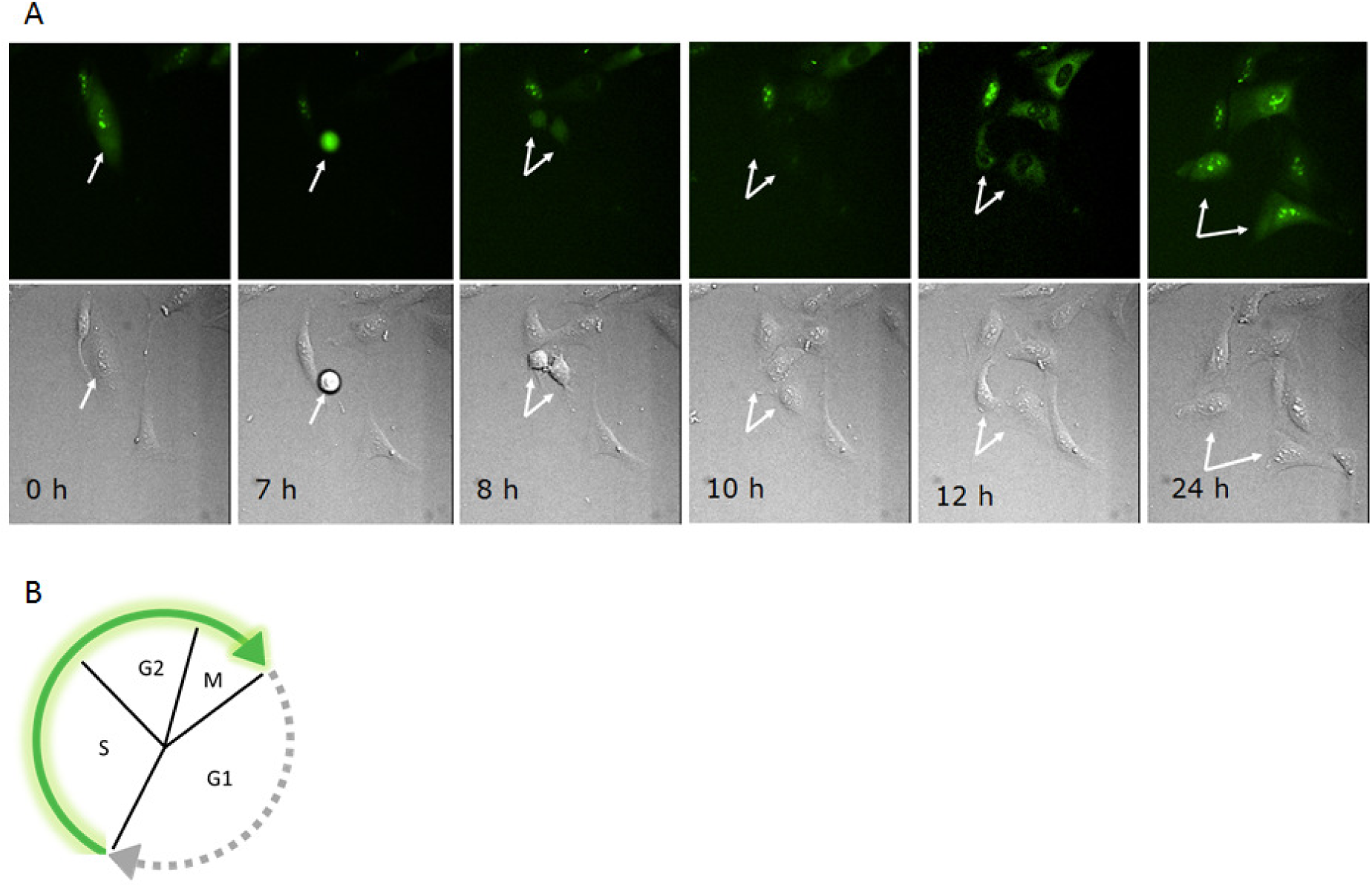
Cas9-GFP-Gemimin fusion protein expression is cell-cycle dependent. (A) Fluorescence images (top) and differential contrast images (bottom) of U2-OS cells expressing Cas9-GFP-Gemimin fusion protein were collected at the indicated time points. The GFP signal was observed to slowly increase throughout the S, G2, and M phases, then rapidly decrease below detection in late mitosis just before cell division, as predicted by the cyclic degradation of geminin alone. This cycling GFP signal suggests that the cycling properties of geminin also apply to this Cas9-GFP-geminin fusion protein. (B) This illustration represents the phases of the cell-cycle in which Cas9-GFP-Gemimin fusion protein is expressed. The GFP signal of the Cas9-GFP-geminin fusion protein is depicted as the thick, green arrow.

### 3.2 Cas9-GFP-geminin increases HDR/NHEJ ratios

We hypothesized that limiting the presence of Cas9 to only the S, G2, and M phases may result in an increased HDR/NHEJ ratio, because HDR is present only in those phases in which geminin is not targeted for degradation. To compare the activities of Cas9-GFP-geminin fusion versus wild-type SpCas9, we chose to compare the activities of both at the endogenous *AAVS1* target site. CEL-I and restriction rragment length polymorphism (RFLP) assays were used to determine the efficiencies of indel formation and ssODN incorporation by both Cas9-GFP-geminin and wild-type SpCas9. Specifically, CEL-I assays were used to indicate the rates of NHEJ repair after cutting by each protein by detecting indels, while the rates of HDR repair were measured via RFLP assay by detecting ssODN incorporation. Figure 3 and 4 indicated that Cas9-GFP-geminin enhanced the HDR/NHEJ ratio significantly in both U2-OS (2.7 fold) and K562 (1.8 fold) cells. Interestingly, our Cas9-GFP-geminin not only decreased NHEJ, but also significantly enhanced HDR. The indels caused by NHEJ may make the original gRNA target sites inaccessible for gRNAs; thus, the decreased NHEJ will likely lead to increased HDR events.

The quantitative results obtained here are comparable to those reported in a paper published by Gutschener, et al., in which a Cas9-geminin fusion protein was reported to increase HDR at the endogenous *MALAT1* locus in HEK293T cells by 1.28-to 1.87-fold compared to wild-type SpCas9, while maintaining similar levels of NHEJ indel formation as with wild-type SpCas9[14]. While Gustchener et al. used plasmid DNA to express their fusion Cas9, another paper by Howden et al. describes delivering another similar geminin-Cas9 fusion protein by means of in vitro transcribed mRNA. In this publication by Howden et al., similar increases in HDR rates were reported while reduced NHEJ levels were also reported, improving the overall HDR/NHEJ ratio in two directions. Howden et al. suggests that this reduction in indels may be due to the transient expression of Cas9 from mRNA versus the likely sustained transcription of Cas9 from the plasmid form[18]. For this reason, we may want to try delivering our fusion protein via in vitro transcribed mRNA in the future.

In summary, we have engineered a Cas9 protein whose expression is cell cycle dependent. This strategy offers a novel alternative approach intended to decrease the NHEJ activity of Cas9, thereby increasing the ratio of HDR to NHEJ and has broader applications than previous small molecule approaches which are often cytotoxic and cell type-dependent. For example, instead of forcing all cells in a population to synchronize, the Cas9-geminin approach synchronizes the enhanced-HDR benefit within each individual cell. While not absolutely required for genome editing applications, the GFP component of our Cas9-GFP-geminin protein provides the first real time observation of spatiotemporal dynamics of this approach and further evidence supporting a positive association between G2/S focused expression and increases in HDR/NHEJ ratios.

**Figure 3.**
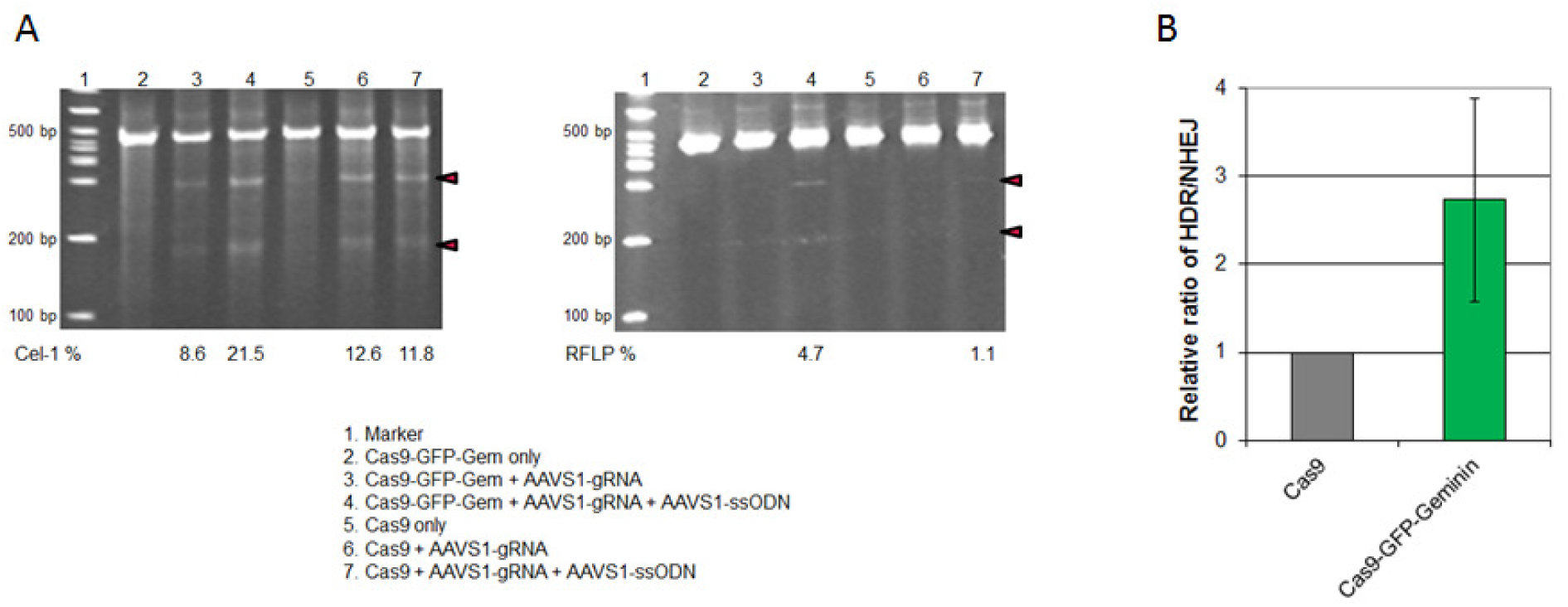
Cas9-GFP-geminin increased HDR/NHEJ ratio in U2-OS cells. (A) The total indels caused by Cas9-GFP-Gem + AAVS1 gRNA (with or without ssODN) or Cas9 + AAVS1 gRNA (with or without ssODN) were measured with Cel-1 nuclease assay, and analyzed using a formula described in **Materials and Methods**. Cas9-GFP-Gem or Cas9 alone was used as negative control. (B) The HDR frequency of Cas9-GFP-Gem + AAVS1 gRNA + ssODN or Cas9 + AAVS1 gRNA + ssODN was measured with RFLP assay (SpeI digestion), and calculated as the ratio of DNA product (indicated with red triangles) to DNA substrate. Cas9-GFP-Gem (with or without gRNA) or Cas9 (with or without gRNA) was used as negative control. (C) This plot depicts the relative ratio of HDR to NHEJ of Cas9-GFP-geminin versus Cas9. An increased HDR/NHEJ ratio is seen in those U2-OS cells transfected with Cas9-GFP-geminin. Standard deviation (error bars) were calculated from three biological replicates.

**Figure 4.**
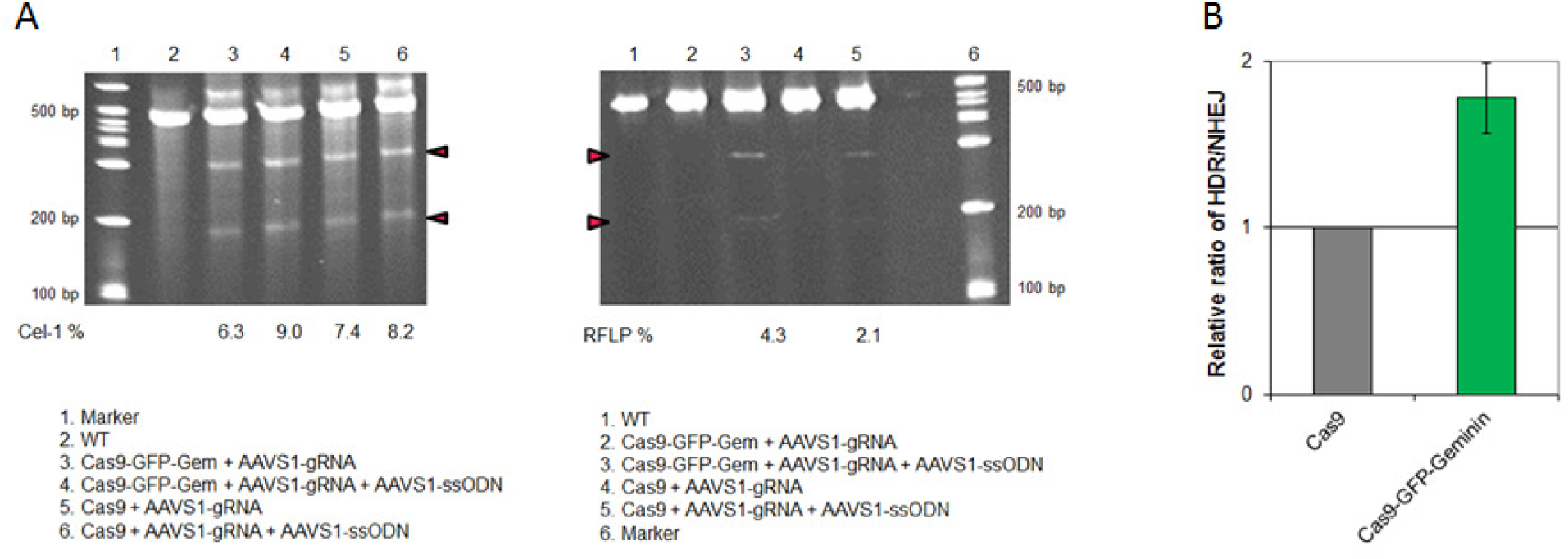
Cas9-GFP-geminin increased HDR/NHEJ ratio in K562 cells. (A) The total indels in K562 cells were measured with Cel-1 nuclease assay. (B) The HDR frequency in K562 cells was measured with RFLP assay (SpeI digestion). (C) This plot depicts the relative ratio of HDR to NHEJ of Cas9-GFP-geminin versus Cas9. An increased HDR/NHEJ ratio is seen in those K562 cells transfected with Cas9-GFP-geminin. Standard deviation (error bars) were calculated from three biological replicates.

## ACKNOWLEDGEMENTS

We thank Patrick Sullivan for R&D administrative support. We thank Jacob Lamberth for technical assistance.

## SUPPLEMENTAL MATERIALS

**Table S1.**
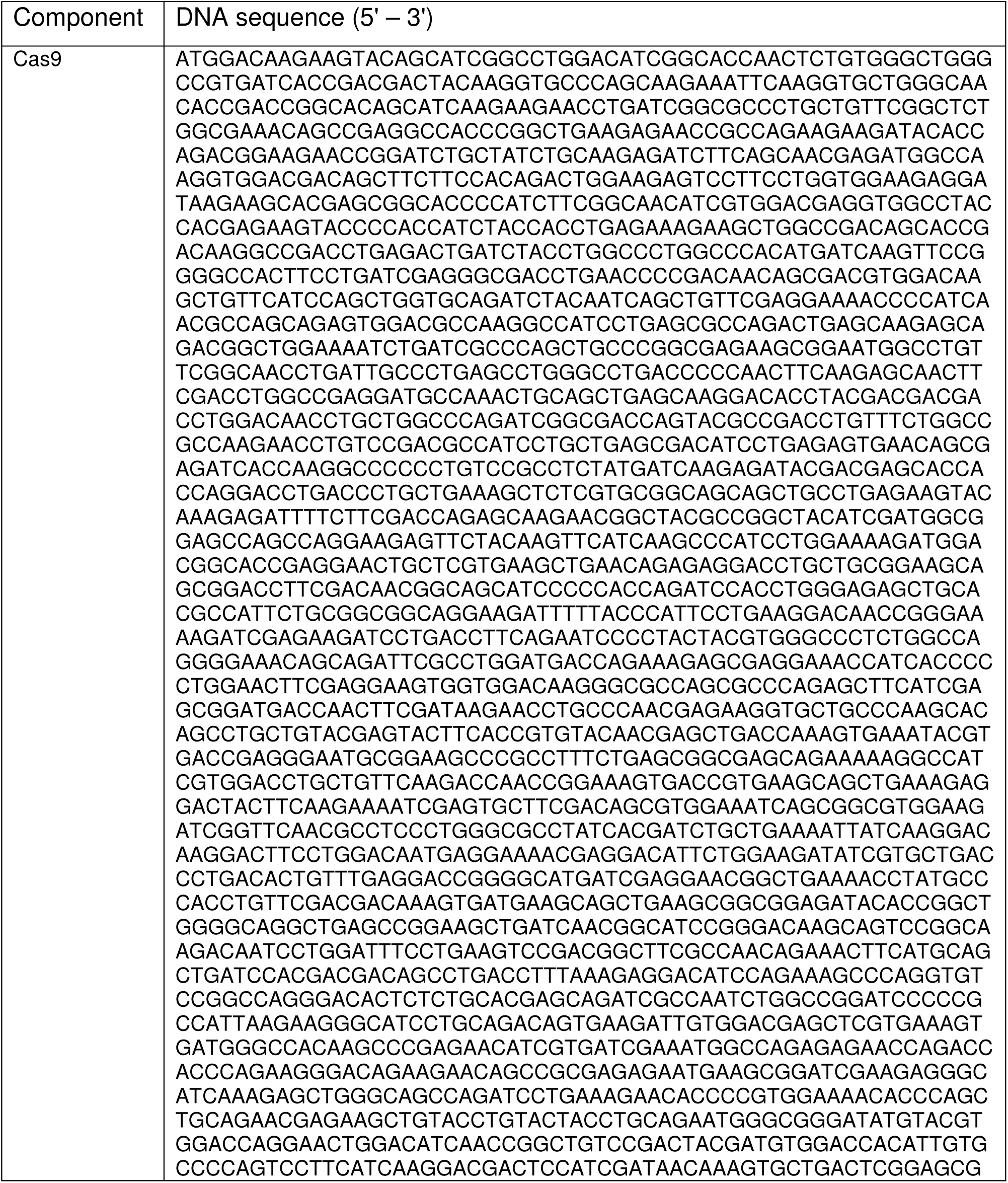
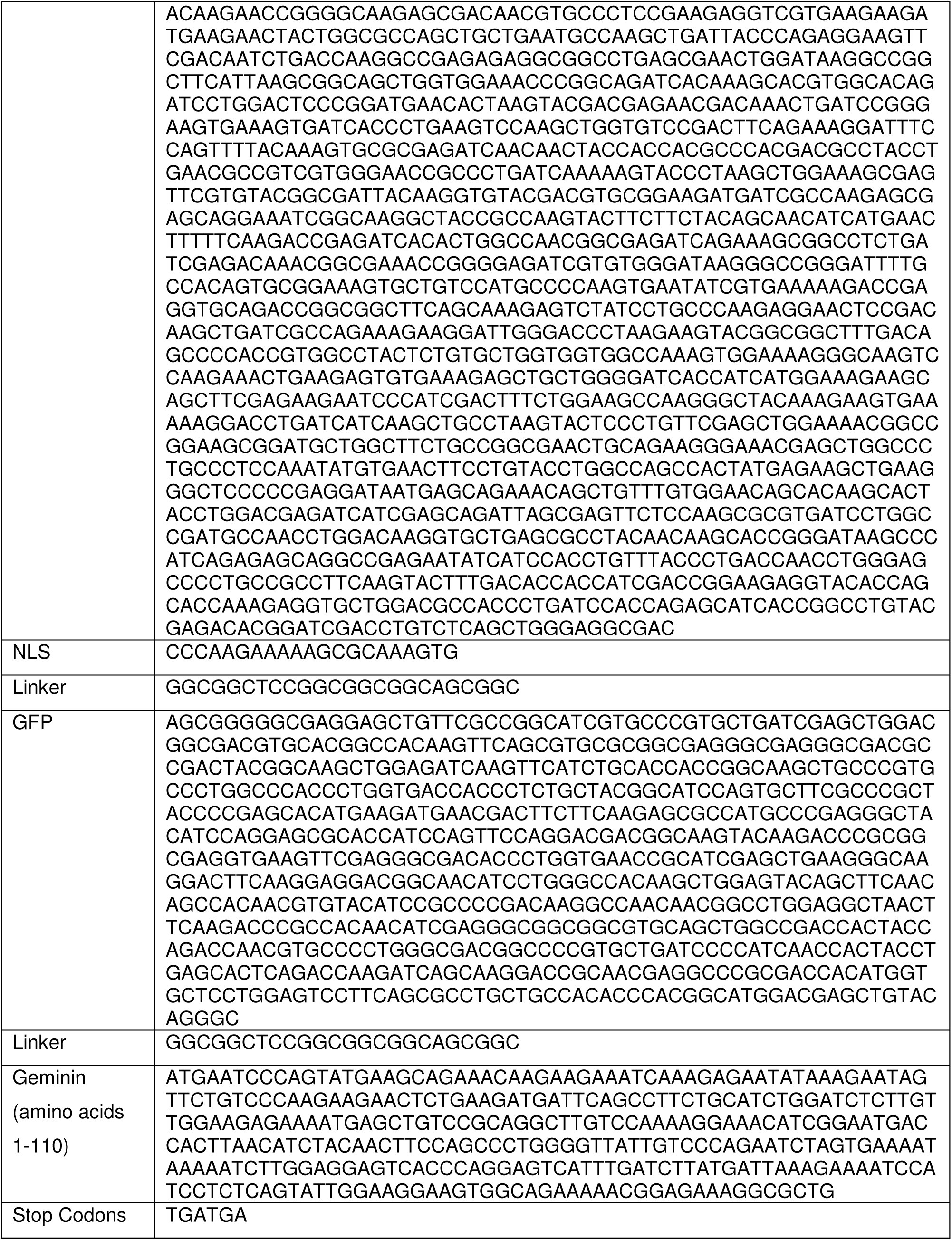
DNA Sequence of Cas9-NLS-GFP-geminin Fusion Protein

**Table S2.**
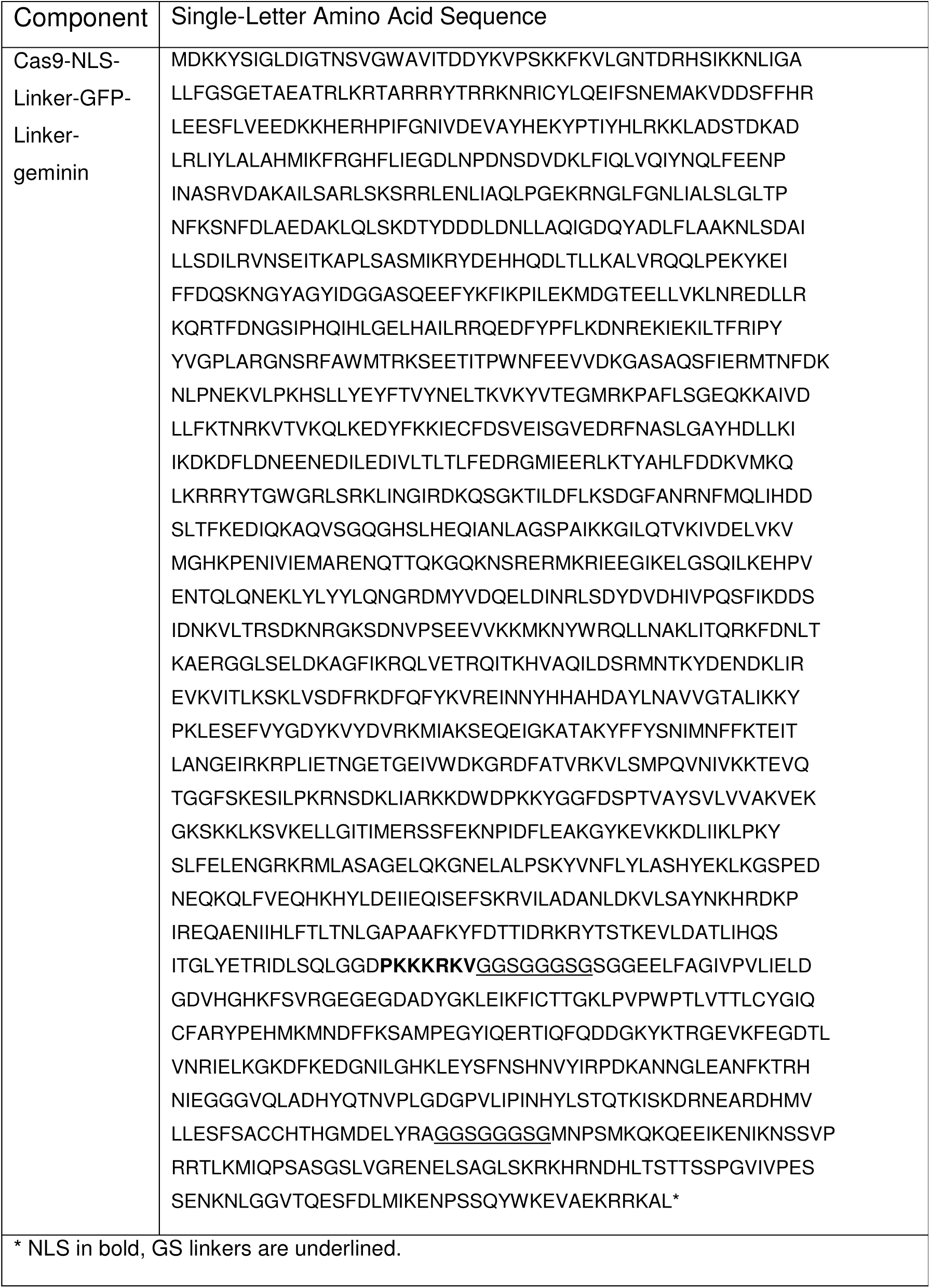
Amino Acid Sequence of Cas9-NLS-GFP-geminin Fusion Protein

